# Gene fractionation and function in the ancient subgenomes of maize

**DOI:** 10.1101/095547

**Authors:** Simon Renny-Byfield, Eli Rodgers-Melnick, Jeffrey Ross-Ibarra

## Abstract

The maize genome experienced an ancient whole genome duplication approximately 10 million years ago and the duplicate subgenomes have since experienced reciprocal gene loss (fractionation) such that many genes have returned to single-copy status. This process has not affected the subgenomes equally; reduced gene expression in one of the subgenomes mitigates the consequences of mutations and gene deletions and is thought to drive higher rates of fractionation. Here we take advantage of published genome-wide SNP and phenotype association data to show that, in accordance with predictions of this model, paralogs with greater expression contribute more to phenotypic variation compared to their lowly expressed counterparts. Furthermore, paralogous genes in the least-fractionated subgenome account for a greater degree of phenotypic diversity than those resident on the more-fractionated subgenome. We also show that the two subgenomes of maize are distinct in epigenetic characteristics. Intriguingly, analysis of singleton genes reveals that these differences persist even after fractionation is complete.

## 1 Introduction

The advent of the plant genomics era has revealed the ubiquity and importance of whole genome duplication (WGD or polyploidy) in the history of land plants Blanc and Wolfe (2004); Bowers *et al.* (2003); Jiao *et al.* (2011); Leitch and Leitch (2008); Paterson *et al.* (2004); Renny-Byfield and Wendel (2014). Analysis of chromosome number, gene copy number and synteny have shown that all land plants are polyploids, and most lineages have a history of repeated WGD in their ancestry Jiao *et al.* (2011); Renny-Byfield and Wendel (2014).

Recently, much effort has been directed towards understanding the fate of duplicate genes following WGD. In spite of multiple rounds of WGD, analysis of a number of plants has demonstrated that the accumulation of gene duplicates is mitigated by a process of gene fractionation, whereby duplicate homoeologous subgenomes reciprocally lose genes such that most genes eventually return to single copy status Freeling *et al.* (2012); Garsmeur *et al.* (2013); Ibarra-Laclette *et al.* (2013); Langham *et al.* (2004); Renny-Byfield *et al.* (2015); Schnable *et al.* (2011); Tang *et al.* (2012); Thomas *et al.* (2006); Woodhouse *et al.* (2010, 2014). Analyses in *Arabidopsis,* Brassicaceae, cotton and *Zea mays* have further revealed that,following WGD, homoeologous subgenomes do not lose genes at an equal rate Renny-Byfield *et al.* (2015); Schnable *et al.* (2011); Thomas *et al.* (2006), and this bias in gene fractionation can persist through multiple nested duplications Woodhouse *et al.* (2014). Genes residing on the most fractionated of the duplicate subgenomes also tend to be more lowly expressed than their duplicate counterparts, perhaps due to differences in transposable element (TE) accumulation and preferential targeting by 24nt-siRNAs Cheng *et al.* (2016); Renny-Byfield *et al.* (2015); Schnable *et al.* (2011); Woodhouse *et al.* (2014). Indeed, TEs are known to increase methylation levels in nearby genes West *et al.* (2014) and epigenetic silencing of such TEs could then impact nearby gene expression via position effect down-regulation Freeling *et al.* (2012); Hollister *et al.* (2011).

Based on these results, Freeling and co-workers Freeling *et al.* (2012) proposed a model in which the most highly expressed gene of a paralog pair produces more protein product and thus contributes more to the required stoichiometric balance and ultimately to phenotype. This model predicts that mutations impacting function would be more strongly selected in the more highly-expressed paralog, while the more lowly-expressed copy is likely to accumulate mutations and may eventually be lost from the genome. Molecular evolutionary analysis of synonymous and nonsynonymous divergence and diversity supports this model, finding stronger evidence of purifying selection in the less-fractionated maize subgenome Pophaly and Tellier (2015).

A number of testable predictions emerge from this model of expression-driven biased fractionation. Firstly, if expression levels differ between subgenomes, there should also be a corresponding difference in overall contribution to phenotypic variation provided by each subgenome. Secondly, variation lowly-expressed paralogous genes should also contribute less to variation in plant phenotype relative to variation at more highly expressed paralogous loci, regardless of subgenome of origin. Finally, the model predicts that we should expect to see other genomic correlates of differential expression between subgenomes, such as epigenetic signatures associated with gene silencing and heterochromatin formation.

Here we test predictions of the Freeling Freeling *et al.* (2012) model using published data on phenotypic associations and functional genomic analyses in maize (*Zea mays* ssp. *mays*). The maize genome underwent its most recent WGD shortly after divergence from *Sorghum,* and its two subgenomes have experienced biased gene fractionation Schnable *et al.* (2011). Genes on the most-fractionated of the subgenomes (maize2) tend to be expressed at a lower level compared with their paralogous copies on the least-fractionated subgenome (maize1; Schnable *et al.* (2011)), and the maize 1 subgenome appears to have experienced stronger purifying selection Pophaly and Tellier (2015). In accordance with a model of expression-driven biased-fractionation, we find that paralogs with higher expression contribute more to phenotypic variation, and that the maize1 subgenome accounts for a greater degree of phenotypic variation than does maize2. Secondly, we reveal that the most-fractionated maize2 subgenome is more often silenced via cytosine methylation, a result that mirrors comparisons of highly-expressed paralogs and their lowly-expressed counterparts across all genes. Finally, our analyses unexpectedly reveal that while singleton genes lacking a paralogous copy show no difference in expression between maize1 and maize2, maize1 singletons nonetheless explain more phenotypic variation. Overall our results provide evidence supporting the Freeling Freeling *et al.* (2012) model, suggesting that epigenetic differences lead to differential expression and divergence in contribution to phenotypic variation, resulting in subgenome-wide differences in purifying selection that likely explain observations of biased fractionation.

## 2 Materials and Methods

### 2.1 SNP datasets, kinship matrices and modelling heritability

We assess how SNPs residing in each of the two subgenomes of maize contribute to phenotypic variability using published phenotypic data from the maize Nested Assoication Mapping (NAM) population Bradbury *et al.* (2007); Bukowski *et al.* (2015); Swarts *et al.* (2014); Wallace *et al.* (2014). Imputed genotype by sequence data Bukowski *et al.* (2015); Swarts *et al.* (2014) were downloaded from `/iplant/home/glaubitz/RareAlleles/genomeAnnos/AHTP/genotypes/ NAM/namrils_projected_hmp31_MAF02mnCnt2500.hmp.txt.gz.` Phenotypic data from Wallace *et al.* 2014 were downloaded from http://journals.plos.org/plosgenetics/article?id=10.1371/journal.pgen.1004845#s5 and consisted of Best Linear Unbiased Predictors for 45 anatomical and physiological traits, including leaf angle, plant and ear height, days to anthesis and ear row number (see Supplementary File 1). We then applied a regional heritability analysis similar to Yang *et al.* (2011), estimating the additive genetic variance explained by kinship matrices created from sets of SNPs representing different loci or regions of the genome.

We compared a first set of SNPs from within whole-genes models in the maize1 subgenome (least fractionated) to a second set from the maize2 subgenome (most fractionated) using previously published subgenome designations Schnable *et al.* (2011), including only those genes retained as gene duplicates in maize1 and maize2. A third dataset, consisting of SNPs not resident in either maize1 or maize2 syntenic paralogs (i.e. the “rest of the genome”,including non-genic regions) was also generated from the same data.

For each of these SNP datasets we generated a square kinship matrix, using the scaled IBS method implemented in TASSEL 5 (https://bitbucket.org/tasseladmin/tassel-5-standalone.git; Bradbury *et al.* (2007)). We used a Residual Maximum Likelihood (REML) model implemented in LDAK Speed *et al.* (2012) to examine heritability for each phenotypic trait.

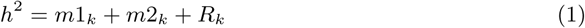

where *m*1_*k*_, *m*2_*k*_ and *R_k_* are kinship matrices for the maize1 subgenome, maize2 subgenome, and the rest of the genome respectively.

We subsequently identified a set of genes that have returned to single copy status following the maize WGD (i.e. genes that have no corresponding duplicate in the alternative subgenome) and separated these genes according to subgenome residency, resulting in a collection of singleton genes for maize1 and maize2. We randomly sampled genes from maize1 to provide the same number of singleton genes observed in maize2. We then generated kinship matrices and estimated the heritability explained for each subsample as described above.

### 2.2 Gene expression analysis

We examined data from the B73 Gene Expression Atlas Sekhon *et al.* (2011) for 70 tissues (available at http://ftp.maizegdb.org/MaizeGDB/FTP/B73%20RNA-SEQ%20Gene%20Atlas/). For each gene we estimated mean expression across 70 tissues and separated each paralog pair into highly-expressed and lowly-expressed categories regardless of the subgenome of origin, provided mean expression was greater than 1.5 fold different between the paralogs (paralog pairs with less than 1.5 fold difference were discarded).

This left us with a collection of two sets of genes of equal size, one set consisting of the most highly expressed of a paralog pair, and the other the more lowly-expressed counterparts. From the remaining genes (i.e. non-paralogous genes), we randomly selected gene pairs, each time segregating genes into lowly- and highly-expressed categories, until we had the same number of differentially expressed pairs (>1.5 fold difference) as that of the paralogous gene sets. We then estimated heritability as described above but using the following model:

*h*^2^ = *pUp_k_* + *pDn_k_* + *rUp_k_* + *rDn_k_* + *R_k_*
where *pUp_k_*, *pDn_k_*, *rUp_k_*, *rDn_k_* and *R_k_* are kinship matrices from highly expressed paralogs, lowly expressed paralogs, highly expressed random genes, lowly expressed random genes, and the remainder of the genome, respectively.

### 2.3 Epigenetic analysis

We evaluated cytosine methylation and chromatin accessibility — as measured by sensitivity to micrococcal nuclease (MNase) — for maize1, maize2, upregulated paralogs, downregulated paralogs, and random sets of upregulated and downregulated genes in 1kb regions surrounding the canonical transcriptional start sites (TSS). DNA methylation data was taken from a whole genome bisulfite sequencing study of B73 Regulski *et al.* (2013). We calculated methylation frequency in 10bp bins, dividing the count of cytosines in the methylated state by the total number of cytosines in each context (CpG, CHG, CHH). We also calculated the frequency of bases in MNase hypersensitive (MNase HS) regions within the same bins, where hypersensitivity was defined according to Rodgers-Melnick *et al.* 2016. We also calculated the frequency of the histone modifications H3K27me2, H3K9me2, and H3K4me3 in maize B73 tissue. H3K9me2 and H3K27me2 CHiP-seq reads were obtained from an earlier study of 1 month-old B73 stalk tissue Gent *et al.* (2014), while H3K4me3 CHiP-seq reads were obtained from a separate study of 14-day B73 whole shoots He *et al.* (2013). For all CHiP-seq reads, we trimmed adaptors, quality-trimmed reads and aligned to the AGPv3 B73 genome according to the methods previously described in Gent *et al.* 2014. For each methylation type, we calculated its frequency in 10bp bins, such that genomic regions overlapping at least 1 read of the given type were considered positive.

## 3 Results

### 3.1 Heritability and the Ancient Subgenomes of Maize

We used published genotypic and phenotypic data from the maize NAM panel Bradbury *et al.* (2007); Bukowski *et al.* (2015); Swarts *et al.* (2014); Wallace *et al.* (2014) to estimate the heritability of 45 phenotypic traits for loci in the two sub-genomes of maize. We first focused analysis on genes retained as syntenic paralogs in both maize1 and maize2. For each of these SNP datasets we used approaches similar to those outlined in Speed *et al.* (2012); Yang *et al.* (2011) to estimate the heritability explained by each set of genes using a residual maximum likelihood model (see methods). Our dataset consisted of 3,195 paralogous genes pairs, with 280,980 and 260,531 SNPs in the maize 1 and maize 2 gene sets, respectively.

Estimates of total heritability for each trait ranged from 6% to 78% (see file S1). For 35 of the 45 traits, loci in the less-fractionated maize1 subgenome explained more heritability than loci in the more-fractionated maize2 sub-genome (binomial test p<0.0002; Fig. 1 and Supplementary File 1). We find identical results restricting the analysis to only traits with high (>40%) heritability (25/30 traits; binomial test, p>0.0004) or commparing mean ranks (Mann-Whitney W = 1543, p-value = 5.9e-07 for all traits, W = 806, p-value = 4.6e-12 high heritability traits, Supplementary Fig. 1).

**Figure 1:**
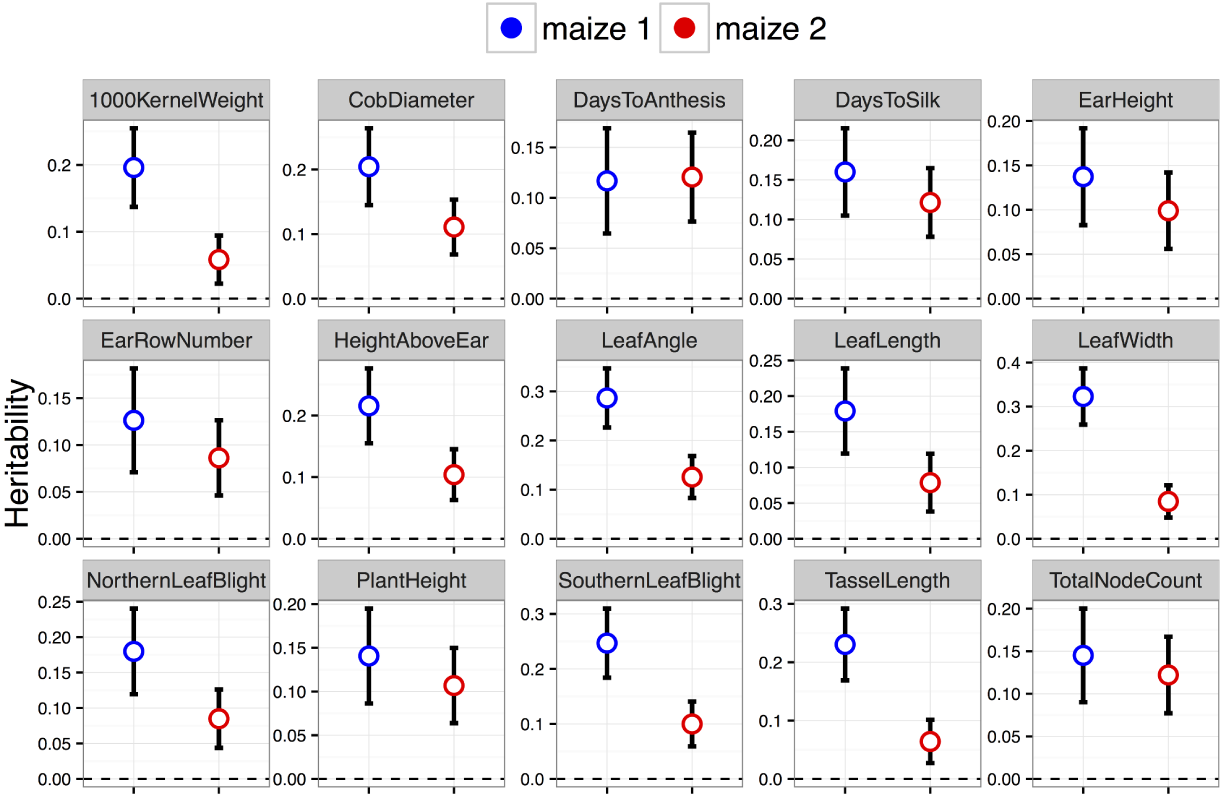
Comparisons of heritability from SNPs in syntenic paralogs from the two subgenomes of maize. Shown are estimates of heritability from 12 representative traits for each of the two subgenomes. Whiskers show 95% confidence intervals generated from the standard error of the mean. Estimates for other traits and for the rest of the genome can be found in the supplement.

We then compared the contributions of singleton genes — those returned to single copy status following maize WGD — from each subgneome (Supplementary Fig. 2 and Supplementary File 1 and 2). Heritability estimates for maize1 and maize 2 singletons varied between 2.6% to 38%. Again maize1 genes explained a greater degree of heritability (25/30 high heritability phenotypes (binomial test p-value <0.0004).

### 3.2 Gene Expression and Heritability

Using RNA-Seq data published as part of the Gene Atlas dataset Sekhon *et al.* (2011), we calculated mean expression for each gene across 70 tissues. We selected pairs of maize1/maize2 paralogs that differed by at least 1.5-fold in expression. We then compared the heritability explained by the group of more highly expressed paralogs to that explained by the group of more lowly-expressed genes; for comparison, we took random pairs of genes and divided them similarly (Fig. 2 and Supplementary File 3). Mean heritability across the 30 high heritability traits differed across the four groups (Kruskal-Wallis test; chi-squared = 27.34, df = 3, p-value <4e-06), with highly expressed paralogs explaining a larger proportion of heritability than their lowly-expressed counterparts (Dunn test, z= -4.5426901, p<0.0005; Supplementary Fig. 3).

**Figure 2:**
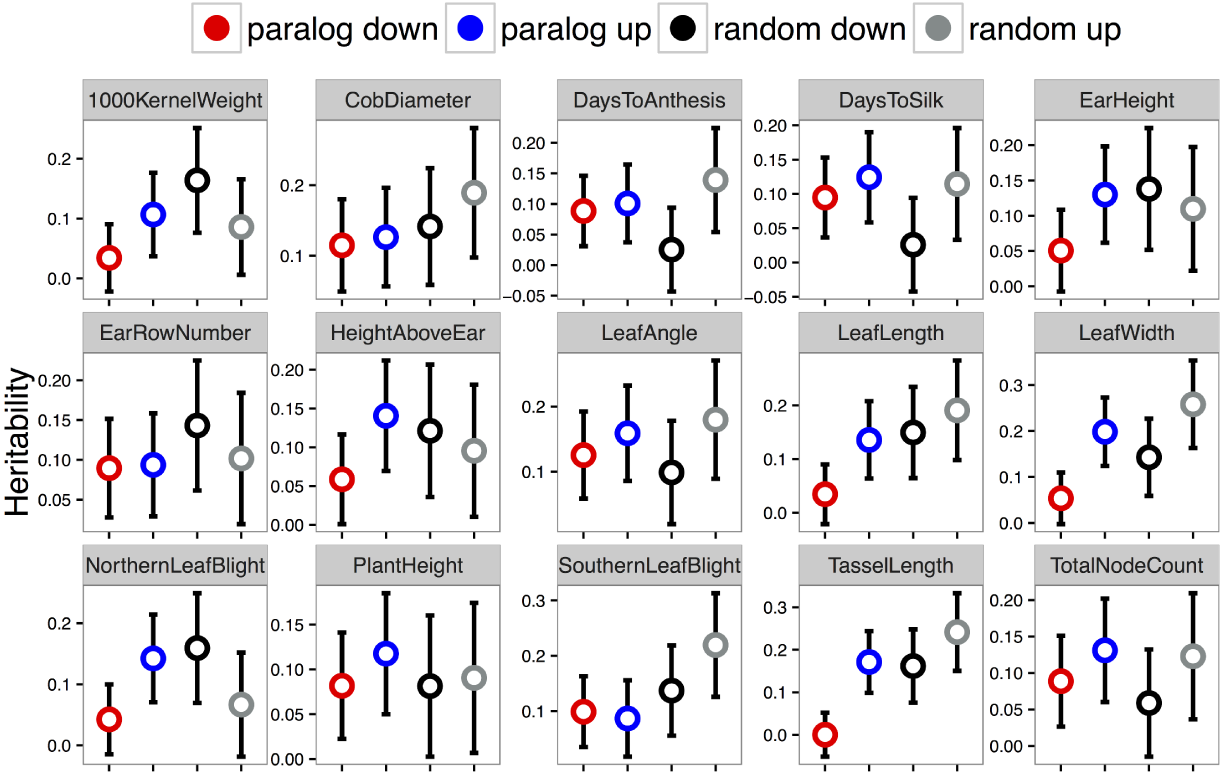
Heritability estimates are shown for a representative set of 12 traits for sets of high and low expressed genes from paralogous pairs as well as randomly selected gene pairs similarly binned into high and low expression. Estimates for other traits and for the rest of the genome can be found in the supplement.

### 3.3 Epigenetic characteristics of maize1 and maize2

We examined previously published epigenetic data Rodgers-Melnick *et al.* (2016) on cytosine methylation, histone modifications, and nucleosome occupancy. We analysed the proportion of methylated sites and compared regions up- and down-stream of the transcription start site (TSS) in genes from maize1 and maize2, revealing that patterns of cytosine methylation (CHH, CHG and CpG) within gene bodies is indistinguishable between each sub-genome (Fig. 3a-c). Ustream of the TSS we observe that maize2 has slightly higher rates of cytosine methylation at CpG and CHG sites (Fig. 3a-b). However, at CHH sites maize1 and maize2 are most noticeably differentiated at ~ 500bp upstream of the TSS (Fig. 3c). Methylation differences between maize1 and maize2 upstream of the TSS remain apparent in singletons as well (Supplementary Fig. 4).

**Figure 3:**
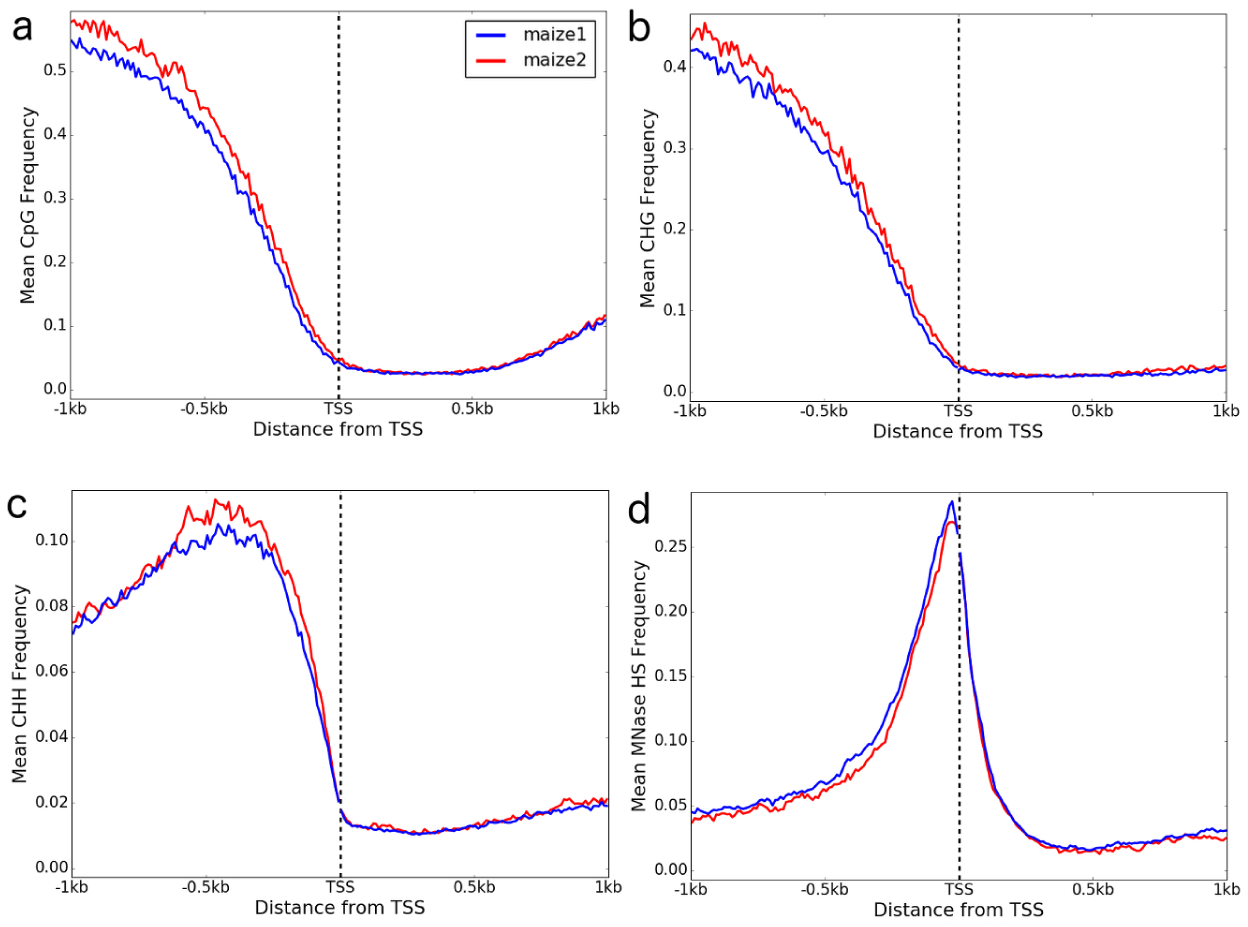
Epigenetic meta-profiles of the two sub-genomes of maize upstream and down-stream of the transcription start site (TSS). Shown are levels of methylation at CpG (a), CHG (b), CHH (c) sites, and chromatin accesibility estimated using MNase (d).

Contrary to cytosine methylation data, Chip-Seq analysis Rodgers-Melnick *et al.* (2016) revealed that methylation rates of h3k27, h3k9 and h3k4 histones does not differ between maize1 and maize2 sub-genomes (Supplementary Figure 5). Analysis of MNase sequencing data Rodgers-Melnick *et al.* (2016) also revealed no difference in chromatin state between maize1 and maize2 within gene bodies, but maize1 has a marginally greater proportion of MNase sensitive regions up-stream of the TSS (Fig. 3d).

### 3.4 Paralogous genes, expression and epigenetic signal

We compared epigenetic signatures around the TSS for 3,195 gene pairs retained as duplicates following an ancient WGD in maize, segregated as before into high and low expression (Fig. 4a-c). Methylation at CHH, CHG and CpG sites is typically higher for the lowly-expressed genes, both upstream of the TSS and inside the gene body when compared with their up-regulatd counterparts (Fig. 4a-c). Randomly paired genes are typically more highly methylated at CHG and CpG sites than paralogous pairs (Fig. 4b & c), an observation also seen in histone methylation patterns (Supplementary Fig 6). In random pairs, while the lowly-expressed gene tends to have higher rates of cytosine methylation (Fig. 4b & c) as seen in paralogous pairs, the pattern is reversed for CHH sites (Fig. 4a).

**Figure 4:**
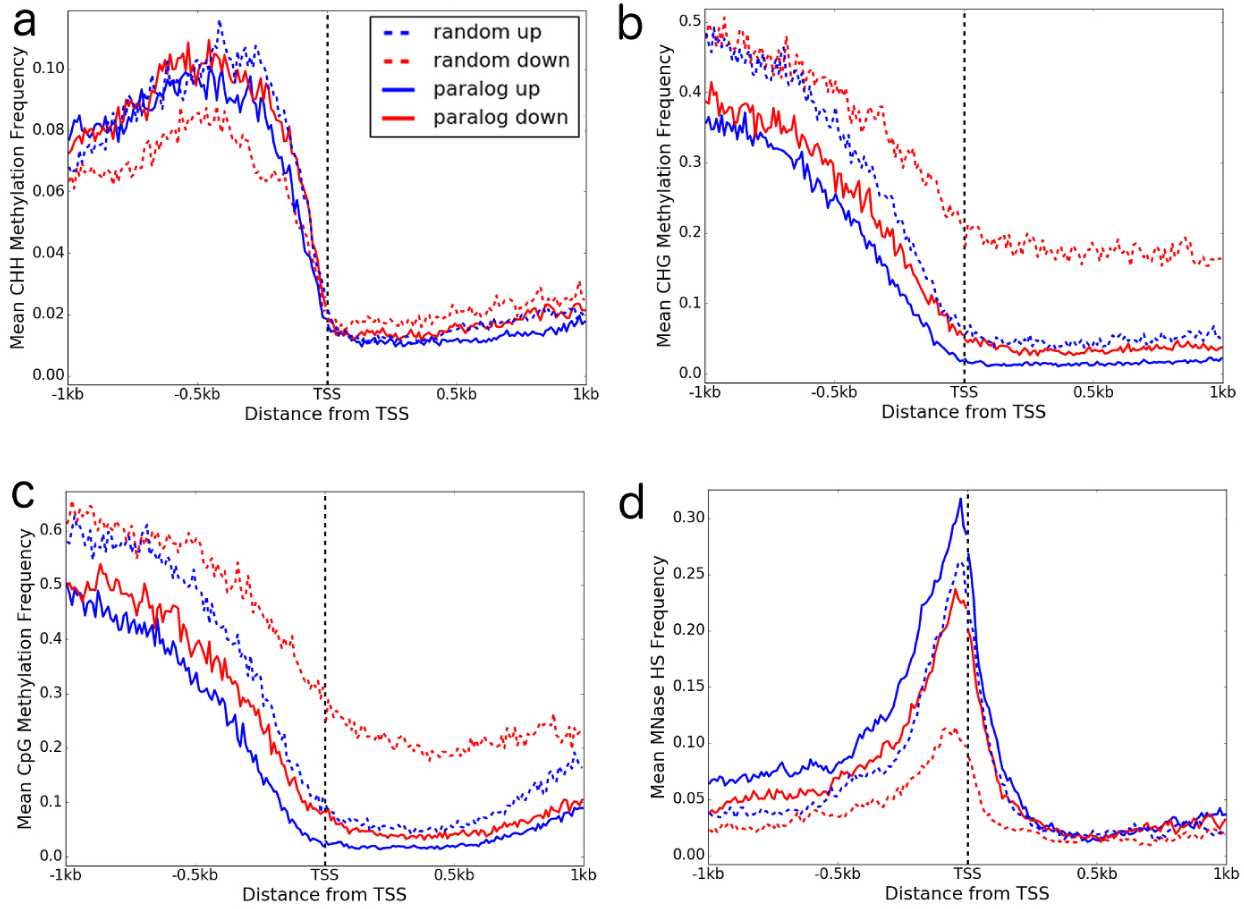
Epigenetic meta-profiles of paralogous genes. For each paralog pair we segregated genes according to expression level (see text). We compared meta-profiles for key epigenetic marks either side of the transcription start site (TSS), including cytosine methlyation rate at CHH (a), CHG (b), and CpG (c) sites, as well as nucleosome association using MNase sensitivity analysis(d).

Finally, we examined nucleosome association with paralog gene expression using MNase sensitivity analysis (Fig. 4d). While all genes tend to have more open chromatin upstream of the TSS, highly expressed genes — in both paralogous and random pairs — appear to have more open chromatin compared to the lowly expressed copy in each pair.

## 4 Discussion

### 4.1 What drives biased fractionation between the ancient sub-genomes of maize?

The evolution of gene content in angiosperm genomes is complicated by the ubiquitous and cyclical nature of whole genome duplication Blanc and Wolfe (2004); Bowers *et al.* (2003); Jiao *et al.* (2011); Leitch and Leitch (2008); Paterson *et al.* (2004); Renny-Byfield and Wendel (2014). Gene loss following WGD, known as fractionation, is of substantial interest due to its impact on the genic content of plant genomes. Researchers have examined what types of genes are retained as duplicates Barker *et al.* (2008); Blanc and Wolfe (2004); Maere *et al.* (2005), what mode of gene deletion prevails Woodhouse *et al.* (2014), and why rates of fractionation vary between duplicate regions of the genome Freeling *et al.* (2012); Garsmeur *et al.* (2013); Renny-Byfield *et al.* (2015); Schnable *et al.* (2011). Following WGD, expression of genes on the most fractionated of the duplicate subgenomes tends be lower than genes on the least-fractionated subgenome. This observation has lead to the suggestion that differences in levels of gene expression might drive differences in rates of gene loss Freeling *et al.* (2012); Schnable *et al.* (2011). Among duplicate gene pairs the gene with the highest level of expression will likely contribute more protein product and mask the impact of SNPs and/or deletions on the more lowly expressed of the gene pair.

In maize, this model of expression-driven biased fractionation predicts that the phenotypic impacts of variants in the most fractionated (maize1) and least fractionated (maize2) sub-genomes differ because maize1 paralogs are typically more highly-expressed relative to maize2. To test this prediction, we estimated the heritability explained by SNPs in maize1, maize2 and separately in up and down-regulated paralogs. We demonstrate that the two sub-genomes of maize have diverged in the degree to which they influence variation in plant phenotype, with paralogous genes on maize1 explaining a greater degree of heritability across the majority of phenotypes examined (Fig. 1). This observation broadly agrees with theory proposed by Freeling and co-workers Freeling *et al.* (2012) and indicates that maize1 is likely more “functional” than the maize2 in terms of its effect on phenotype as well as contribution to gene expression. As selection acts on phenotypic variability, these results are also consistent with the observation that maize1 has been under greater purifying selection since the WGD event Pophaly and Tellier (2015).

We then assessed whether differences in phenotypic variability in maize1 and maize2 are driven specifically by expression differences, rather than some other aspect of subgenome biology. We show that more highly expressed genes, regardless of subgenome of origin, typically explain more variation in phenotype than do their lowly expressed paralogous counterparts (Fig.2). The same pattern is also observed for random sets of non-paralogous genes (Fig.2), suggesting that observed differences in phenotypic variance explained between the subgenomes were likely driven by differences in expression.

At face value, however, this model is contradicted by our observation that heritability differences between subgenomes persist even among loci where fractionation is complete (Supplementary Fig. 2) even though such singletons show no differences in expression (Supplementary Fig. 7). While we do not as yet have a satisfactory explanation for why these differences remain in singletons, we note that as long as the expression level of the (missing) paralog of these singletons was lower than the extant singleton, these results would still be consistent with expression acting as a main driver of phenotypic variation. One alternative explanation is that singleton genes from maize2 could be enriched for psuedogenes or TE derived sequences that have been incorrectly annotated. Inclusion of such nonfunctional loci in the maize2 singleton set might explain their limited contribution to heritability. We briefly examined this possibility by compairng the proportion of single exon genes and the number of exons in maize1 and maize2 singleton genes, but see no differences between the subgenomes (Supplementary Fig. 8).

### 4.2 Epigenetic signals differentiate maize1 and maize2

Expression level of duplicate genes may determine which of the duplicates is deleted, but what might cause expression level differences between subgenomes? Cases of WGD that subsequently result in biased fractionation are thought to be derived from allopolyploid (interspecific hybridisation followed by whole genome duplication) events, whereas those cases where gene loss is equal between the duplicate subgenomes are thought to be due to autopolyplody Garsmeur *et al.* (2013). If the parents of an allopolyploid differ in transposable element (TE) content, fractionation could be triggered by position-effect down-regulation of adjacent genes mediated by TEs Freeling *et al.* (2012); Hollister *et al.* (2011). Indeed, in other plants genes resident in the most fractionated portions of the genome have been shown to exhibit a greater proportion of TEs upstream of the TSS Renny-Byfield *et al.* (2015) and are preferentially targeted by 24nt siRNAs Renny-Byfield *et al.* (2015); Woodhouse *et al.* (2014). Under this model, the fate of duplicate genes would be set by the TEs resident in each ancestral genome, driving future biases in gene loss once united in a single allopolyploid nucleus.

If such early variability in TE content drives expression differences between subgenomes, we might expect this to be reflected at the epigenetic level. In this study, we show that maize1 and maize2 are differentiated in a number of epigenetic characteristics (Fig. 3a-c), most notably upstream of the TSS. Though not highlighted previously, earlier analyses identify marginal differences similar to those reported here West *et al.* (2014). Methylation of DNA is generally an inhibitor of gene expression and TEs, and the pattern we observe for maize1 and maize2 is consistent with observations regarding gene expression Renny-Byfield *et al.* (2015); Schnable *et al.* (2011); Woodhouse *et al.* (2010, 2014) and TE content Renny-Byfield *et al.* (2015); ? from a number of plant species. Not only are sub-genomes differentiated via cytosine methylation, but also in MNase sensitivity (Fig. 3d). Although the signal is weak, there is a consistently higher frequency of MNase sensitive nucleotides in the maize1 subgenome upstream of the TSS, suggesting a more open chromatin environment. In individual paralog pairs, the more highly expressed gene typically exhibits a lower proportion of methylated cytosines up-stream of the TSS and in gene bodies(Fig. 4a-c), as well more open chromatin (Fig. 4d), consistent with a model of gene silencing via methylation. Together, these data support our hypothesis that expression differences between subgenomes may derive from epigenetic differentiation.

## 5 Supplementary Material

Supplementary tables and figures are available online ().

## 6 Acknowledgements

We thank the National Science Foundation Division of Integrative Organismal Systems (grant number 1238014) for funding and Jeff Glaubitz, Edward Buckler and Michelle Stitzer for constructive comments on the manuscript.

